# ENVIRONMENTS and EOL: identification of Environment Ontology terms in text and the annotation of the Encyclopedia of Life

**DOI:** 10.1101/011403

**Authors:** Evangelos Pafilis, Sune P Frankild, Julia Schnetzer, Lucia Fanini, Sarah Faulwetter, Christina Pavloudi, Aikaterini Vasileiadou, Patrick Leary, Jennifer Hammock, Katja Schulz, Cynthia Sims Parr, Christos Arvanitidis, Lars Juhl Jensen

## Abstract

**Summary:** The association of organisms to their environments is a key issue in exploring biodiversity patterns. This knowledge has traditionally been scattered, but textual descriptions of taxa and their habitats are now being consolidated in centralized resources. However, structured annotations are needed to facilitate large-scale analyses. Therefore, we developed ENVIRONMENTS, a fast dictionary-based tagger capable of identifying Environment Ontology (ENVO) terms in text. We evaluate the accuracy of the tagger on a new manually curated corpus of 600 Encyclopedia Of Life (EOL) species pages. We use the tagger to associate taxa with environments by tagging EOL text content monthly, and integrate the results into the EOL to disseminate them to a broad audience of users.

**Availability and implementation:** The software and the corpus are available under the open-source BSD and the CC-BY-NC-SA 3.0 licenses, respectively, at http://environments.hcmr.gr

## 1 INTRODUCTION

The Encyclopedia of Life (EOL; http://eol.org/) is a web resource offering biodiversity knowledge summaries of the world’s species to a vast audience (Parr *et al*., 2014). It currently aggregates content from more than 250 providers. These include textual descriptions about the biology, such as habitat, of more than 900,000 taxa.

The Environment Ontology (ENVO) project aims to provide a controlled, structured vocabulary to support annotation of organisms with environmental descriptors (Buttigieg *et al*., 2013). The ontology comprises ∼1600 terms and is part of recommended (meta-)genomic metadata standards (Yilmaz *et al*., 2011). Having the environmental information contained in EOL annotated in the form of ENVO terms, rather than as free text, would enhance search capabilities and enable users to easily compile summary statistics on, for example, the ecological distribution of any taxa. However, manually annotating all EOL entries with ENVO terms would be highly time demanding. An attractive alternative is to use text mining to automatically tag ENVO terms. So far, the few efforts to perform named entity recognition of environments have focused on tagging bacteria biotopes (Bossy *et al*., 2013) or made use of generic tools, not optimized for the task (Thessen and Parr, 2014).

Here we present ENVIRONMENTS, a tagger capable of identifying ENVO terms with sufficient accuracy and speed to be useful for annotating large text corpora. To benchmark the method, we developed a new gold standard corpus of manually annotated EOL taxon pages. Last but not least, we have extended the EOL web resource with ENVO terms for each taxon, which are automatically mined from their textual descriptions.

## 2 ENVIRONMENT ONTOLOGY TERM TAGGER

ENVIRONMENTS identifies ENVO terms in text using the same fast dictionary-based tagging engine as in Pafilis *et al.*, 2013. The command-line tool requires only a single parameter, namely the path to a folder with the text files to be processed.

We constructed a dictionary based on ENVO by extracting all names and synonyms from the OBO file, excluding broad synonyms, obsolete terms, terms describing foods rather than environments, and terms representing organisms and tissues, which are better captured by other ontologies. Names connected to multiple terms were assigned to the one that best captures its meaning, by ranking the terms based on whether the name was the primary name, an exact synonym, a narrow synonym, or a related synonym of the term. Because ENVO usually lists only the singular noun forms, we automatically generated plural and adjective forms.

A small fraction of the names will result in many false positives due to homonymy. We created a block list of such names by inspecting text for all names that appeared more than 2000 times in Medline and EOL. To find important synonyms missing in ENVO, we tagged the habitat and ecology sections of 1,342,968 EOL pages that had even (not odd) identifiers, and inspected all words occurring more than 100 times in untagged text segments. Based on this analysis and false negatives found in the development part of the curated corpus, we added 142 synonyms to the dictionary.

## 3 MANUALLY CURATED CORPUS

To construct an evaluation corpus that covers diverse environment types, we retrieved 313,269 EOL species pages representing the clades Actinopterygii, Annelida, Arthropoda, Aves, Chlorophyta, Mammalia, Mollusca, Streptophyta. From these we extracted the sections about behavior, biology, dispersal, distribution, ecology, habitat, legislation, migration, reproduction, and trophic strategy. After discarding very short (<100 words) and very long (>1,000 words) pages, we randomly selected 75 species from each of the eight clades. The resulting 600 species pages were randomly distributed among six annotators; 20% of the pages were given to two annotators to allow for assessment of inter-annotator agreement (IAA). Each person independently annotated ENVO terms in the text, with no knowledge of which pages had been assigned to a second annotator. Based on the shared abstracts we find that the median pairwise Cohen’s kappa is 0.65, implying that the overall IAA is acceptable despite the difficulty of the annotation task (a Cohen's kappa value of 0 indicates random agreement, while a value of 1 total agreement). The corpus was partitioned so that pages with even and odd EOL identifiers were used for development and final evaluation, respectively.

## 4 PERFORMANCE EVALUATION

Since the ENVIRONMENTS tagger recognizes names within text and links them to ENVO terms, we benchmarked both aspects of its performance. To quantify to which extent the tagger recognizes the same text fragments as the annotators, we calculated precision and recall at the mention level, considering both exact and partially overlapping matches as true positives. On the evaluation part of the corpus, this resulted in 87.8% precision and 77.0% recall, corresponding to an F1 score of 82.0%. For the matches that were considered true positive for the recognition task, we further evaluated if the tagger linked them to the same ENVO terms as the annotators did. In 87.1% of cases, the tagger and the annotator agreed on at least one ENVO term.

## 5 ANNOTATION OF ENCYCLOPEDIA OF LIFE

To realize the full potential of any text-mining system, it is important that it is adopted by the broad community and that its results are disseminated to the intended end users. To this end, we have integrated the ENVIRONMENTS tagger with the EOL web resource to provide users with ENVO terms for each taxon. Each month, we rerun the tagger on all English text in environment-related sections of EOL. As of October 2014, this gave rise to 1,077,522 annotations of ENVO terms for 234,582 EOL taxa.

We make these environment annotations available to end users in three different ways. First, we show them within the EOL taxon web pages, which provide links to the relevant paragraphs in the textual descriptions for each ENVO term (Figure 1). Second, they can be queried through the web interface or the application programming interface of the new EOL/Traitbank semantic web data repository of organismal traits (http://eol.org/traitbank). Third, the full annotation dataset can be downloaded in tab-delimited format from http://downloads.jensenlab.org/EOL.

**Figure 1.**
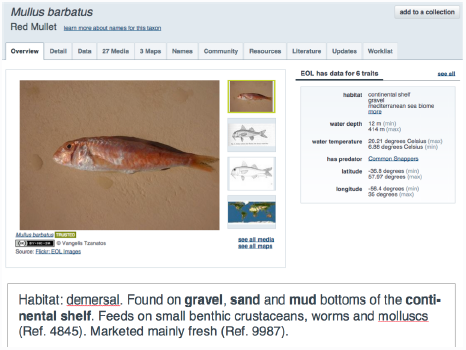
*Top*: The “Overview” tab of the EOL taxon pages show a subset of the ENVO terms obtained through text mining; an extended list of such terms is available in the “Data” tab. *Bottom*: The latter list provides links to the EOL text sections where each term was found (highlighted in bold).

## 6 FUTURE WORK

The ENVIRONMENTS tagger is applicable to other large sources of text than EOL. For example, it can be applied to text fields, such as isolation source, in Genbank (Hirschman *et al*., 2008). Combined with the SPECIES tagger (Pafilis *et al*., 2013), it can also be used to extract species–environment pairs from the scientific literature, like legacy biodiversity literature (Gwinn and Rinaldo, 2009).

## ACKNOWLEDGEMENTS

### Funding

The Encyclopedia Of Life Rubenstein Fellows Program [CRDF EOL-33066-13/E33066], the LifeWatchGreece Research Infrastructure [384676-94/GSRT/NSRF(C&E)], and the Novo Nordisk Foundation Center for Protein Research [NNF14CC0001].

### Conflict of Interest

none declared.

